## Supplementary Information (Supp. Tables 1-2 + Supp. Figs. 1-9) for "CO-OPTION OF NECK MUSCLES SUPPORTED THE VERTEBRATE WATER-TO-LAND TRANSITION"


Eglantine Heude[1,2,*], Hugo Dutel[3,4,5], Frida Sanchez-Garrido[1], Karin D. Prummel[6,7], Robert Lalonde[7], France Lam[8], Christian Mosimann[6,7], Anthony Herrel[9,10,11,12], Shahragim Tajbakhsh[13,14]

[1]PHYMA, Département Adaptations du Vivant, Muséum national d'Histoire naturelle, CNRS UMR 7221; Paris, 75005, France.
[2]Institut de Génomique Fonctionnelle de Lyon, École Normale Supérieure de Lyon, CNRS UMR 5242, Université Claude Bernard Lyon-1; Lyon, 69007, France.
[3]Bristol Palaeobiology Research Group, School of Earth Sciences, University of Bristol; Bristol, BS8 1RJ, United Kingdoms.
[4]Université de Bordeaux, CNRS, MCC, PACEA, UMR 5199; Pessac, 33600, France.
[5]Craniofacial Growth and Form, Hôpital Necker - Enfants Malades, Paris, 75015, France
[6]Department of Molecular Life Sciences, University of Zurich; Zurich, CH-8057, Switzerland.
[7]Department of Pediatrics, Section of Developmental Biology, University of Colorado School of Medicine, Anschutz Medical Campus; Aurora, CO 80045, USA.
[8]Core Facilities - Institut de Biologie Paris Seine (IBPS), Sorbonne Universités; Paris, 75005, France.
[9]MECADEV, Département Adaptations du Vivant, Muséum national d'Histoire naturelle, CNRS UMR 7179; Paris, 75005, France.
[10]Department of Biology, Evolutionary Morphology of Vertebrates, Ghent University; Ghent, 9000, Belgium.
[11]Department of Biology, University of Antwerp; Wilrijk, 2610, Belgium.
[12]Naturhistorisches Museum Bern; Bern, 3005, Switzerland.
[13]Department of Developmental & Stem Cell Biology, Stem Cells & Development Unit, Institut Pasteur, Université Paris Cité; Paris, 75015, France.
[14]CNRS UMR3738, Institut Pasteur; Paris, 75015, France.

* To whom correspondence should be addressed:
Dr. Eglantine Heude
Institut de Génomique Fonctionnelle de Lyon
École Normale Supérieure de Lyon
CNRS UMR 5242, Université Claude Bernard Lyon-1
Lyon, 69007, France.


### SUPPLEMENTARY TABLES

**Supplementary Table 1.** Parameters used for µCT acquisitions at the XTM Facility, Palaeobiology Research Group, University of Bristol.

| Genus | *Ambystoma* | *Anolis* | *Protopterus* |
|---|---|---|---|
| Scanner | Nikon XTH 225ST | Nikon XTH 225ST | Nikon XTH 225ST |
| Voxel size (mm) | 0.0077 | 0.01982 | 0.01144 |
| Voltage (kV) | 80 | 85 | 90 |
| Current (µA) | 96 | 140 | 145 |
| Exposure time (ms) | 500 | 500 | 500 |
| Projections | 3141 | 3141 | 3141 |
| Contrast agent | 2.5% PMA | 5% PMA | 5% PMA |

**Supplementary Table 2.** Parameters used for PPC-SRµCT acquisitions on BM19 at the European Synchrotron Facility.

| Genus | *Latimeria* (ZSM 28409) | *Polypterus* / *Tylototriton* |
|---|---|---|
| Voxel size (mm) | 0.03045 | 0.00655 |
| Average energy (keV) | 63.2 | 33.8 |
| Optics | Hasselbald cinema optic | Tandem Hasselblad 2x |
| Filter | Al 2<br>Cu 0.25<br>W 0.25 | Al 2<br>Cu 0.1 |
| Propagation distance (mm) | 2800 | 3000 |
| Sensor | FReLoN 2K14 | FReLoN 2K14 (Frame transfer mode) |
| Scintillator | LuAG:Ce 750 µm | LuAG:Ce 250 µm |
| Insertion device | W150 | U17.6 second harmonic |
| ID Gap (mm) | 70 | 14 |
| Scan geometry | 360, half-acquisition, vertical series 7 mm | 360, half-acquisition, vertical series 5 mm |
| Exposure time (ms) | 100 | 200 |
| Projections | 4998 | 3000 |
| Time per scan (min) | 9.8 | 12 |
| Reconstruction | Phase retrieval | Single distance phase retrieval, 16 bits conversion, vertical concatenation, ring artefacts correction |

### SUPPLEMENTARY FIGURE LEGENDS

**Supplementary Fig. 1. Cardiopharyngeal mesoderm contribution to the cucullaris muscle in zebrafish larvae.**

3D rendering of whole-mount immunofluorescent staining showing *tbx1*-lineage reporter expression (GFP, green) in head/trunk-connecting muscles (MyHC, magenta) (a-c) of the zebrafish larva #1 shown in (Fig. 2a-i) 5 days post-fertilization (dpf). The data reveal high recombination efficiency in the heart and branchial arch musculature including the cucullaris (yellow arrowheads, a-b) and coracobranchial (c) myofibers. In contrast, the GFP is not detected in somitic-derived pectoral fin, ventral hypaxial and sternohyoid myofibers (a-c) and no recombination is observed in inducible CreERT2 reporter larvae without OHT treatment (d-e), demonstrating Cre-mediated recombination specificity. The OHT treatments in inducible *tbx1*-lineage reporter larvae resulted in reproducible recombination and redundant GFP expression in the cucullaris as shown in specimens #2 to #5 (f-s).

Abbreviations: bam, branchial arch musculature; cbm, coracobranchial muscle; dpf, days post-fertilization; e, eye; h, heart; pfm, pectoral fin musculature; sh, sternohyoid muscle; vhp, ventral hypaxial muscles; y, yolk. Scale bar in (j), for (a, d-g) 200 µm, for (b-c, h-s) 50 µm.

**Supplementary Fig. 2. Neuromuscular system at the head trunk transition of zebrafish larvae.** Whole-mount immunofluorescent stainings of the muscular (MyHC, magenta) and nervous systems (Ac-Tub, yellow) of zebrafish larvae 5- and 7-days post-fertilization (dpf) acquired by light sheet fluorescent microscopy. Maximum intensity projections (a, e, i) and lateral 3D renderings of the neuromuscular system (b-d, f-h, j-l) at the level indicated on schemes, with muscular and nervous structures separated in (c-d, g-h, k-l). See also Supplementary Movies 1 and 2 for interactive details.
bam, branchial arch musculature; dhp, dorsal hypaxial musculature; dpf, days post-fertilization; e, eye; eom, extraocular muscles; ep, epaxial musculature; fc, facial cranial nerve VII; h, heart; hg, hypoglossal cranial nerve XII; ltl, lateral line; mdm, mandibular muscles; of, olfactory bulb; pfm, pectoral fin musculature; sh, sternohyoid muscle; sp, spinal nerves; tg, trigeminal cranial nerve V; vg, vagus cranial nerve X; vhp, ventral hypaxial musculature. Scale bar in (d), for (a-h) 200 µm, for (i-l) 50 µm.

**Supplementary Fig. 3. Reconstructions of the musculoskeletal system at the head/trunk transition of a juvenile zebrafish.**
(a) More 3D-renderings corresponding to data presented in (Fig. 3a). (b) Preotic and postotic virtual frontal sections from CT scan raw data showing the segmented structures of interest.

**Supplementary Fig. 4. Reconstructions of the musculoskeletal system at the head/trunk transition of a juvenile bichir.**
(a) More 3D-renderings corresponding to data presented in (Fig. 3b) with vagus nerve (nX) reconstruction (in yellow). (b) Preotic and postotic virtual frontal sections from CT scan raw data showing the segmented structures of interest.

**Supplementary Fig. 5. Reconstructions of the musculoskeletal system at the head/trunk transition of a juvenile coelacanth.**
(a) 3D-renderings of the somitic- and cardiopharyngeal-derived musculature with vagus nerve (nX) reconstruction (in yellow). The posterior branchial musculature is shown in light blue and the putative cucullaris described in Sefton *et al.* (2018) and Johnson (2022) is shown in light grey. Our analysis supports the absence of cucullaris homologous muscle in coelacanth. (b) Preotic and postotic virtual frontal sections from CT scan raw data showing the segmented structures of interest.

**Supplementary Fig. 6. Reconstructions of the musculoskeletal system at the head/trunk transition of a juvenile lungfish.**
(a) More 3D-renderings corresponding to data presented in (Fig. 3c) with vagus nerve (nX) reconstruction (in yellow). (b) Preotic and postotic virtual frontal sections from CT scan raw data showing the segmented structures of interest.

**Supplementary Fig. 7. Reconstructions of the musculoskeletal system at the head/trunk transition of a juvenile axolotl.**
(a) More 3D-renderings corresponding to data presented in (Fig. 3d). (b) Preotic and postotic virtual frontal sections from CT scan raw data showing the segmented structures of interest.

**Supplementary Fig. 8. Reconstructions of the musculoskeletal system at the head/trunk transition of a juvenile newt.**
(a) More 3D-renderings corresponding to data presented in (Fig. 3e). (b) Preotic and postotic virtual frontal sections from CT scan raw data showing the segmented structures of interest.

**Supplementary Fig. 9. Reconstructions of the musculoskeletal system at the head/trunk transition of a juvenile lizard.**
(a) More 3D-renderings corresponding to data presented in (Fig. 3f). (b) Preotic and postotic virtual frontal sections from CT scan raw data showing the segmented structures of interest.

### SUPPLEMENTARY FIGURES

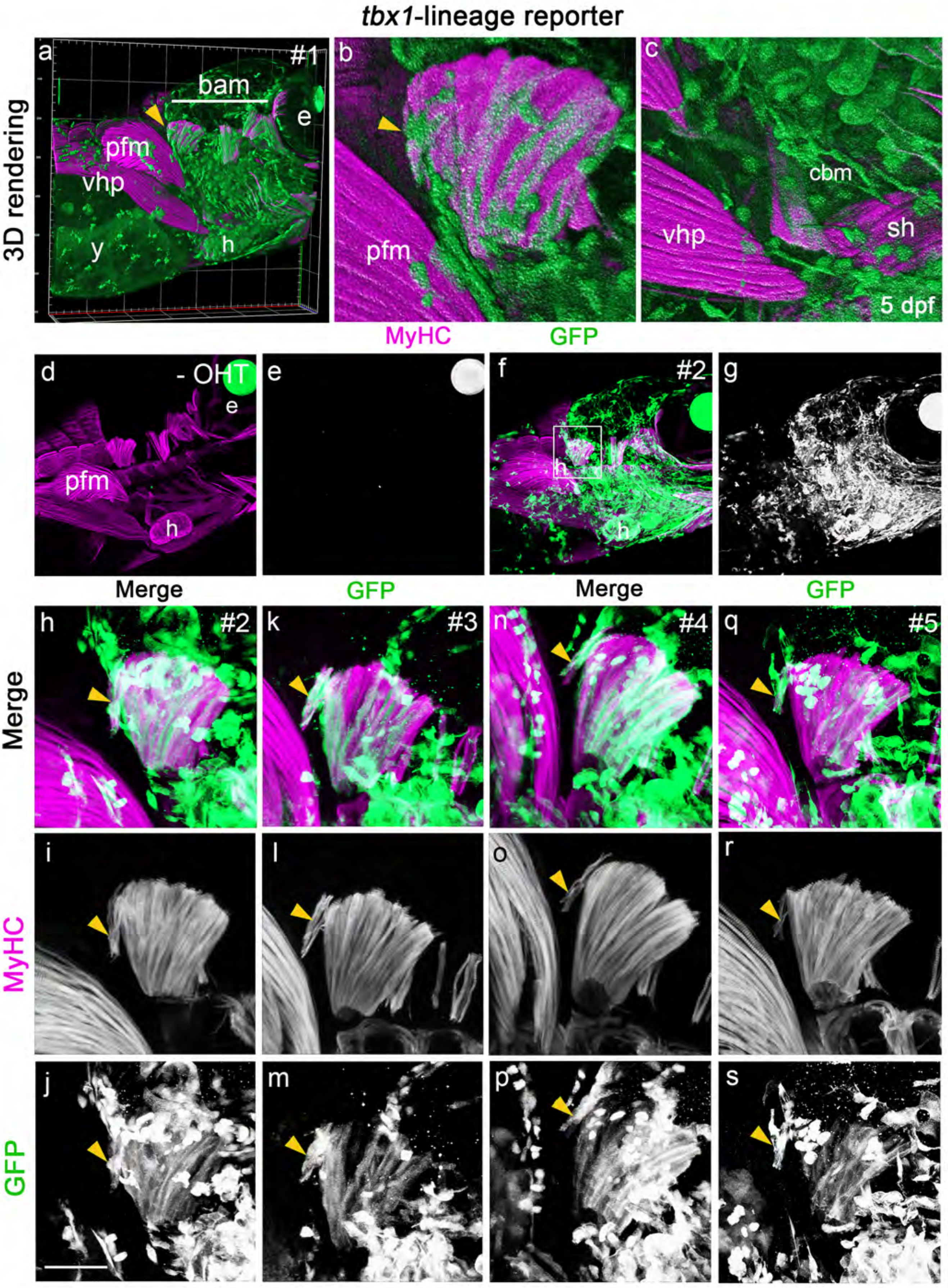


Supplementary Fig. 1

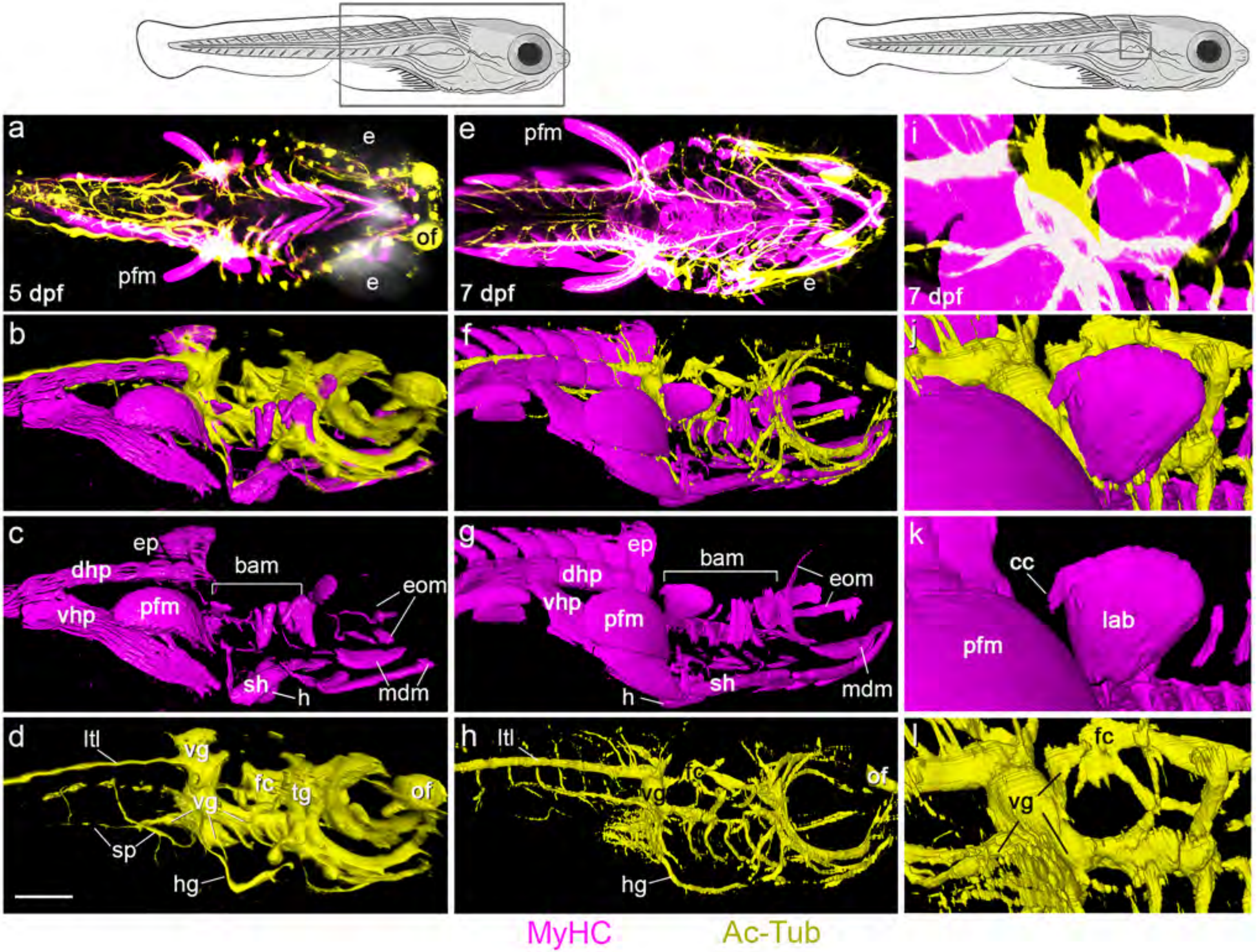


Supplementary Fig. 2

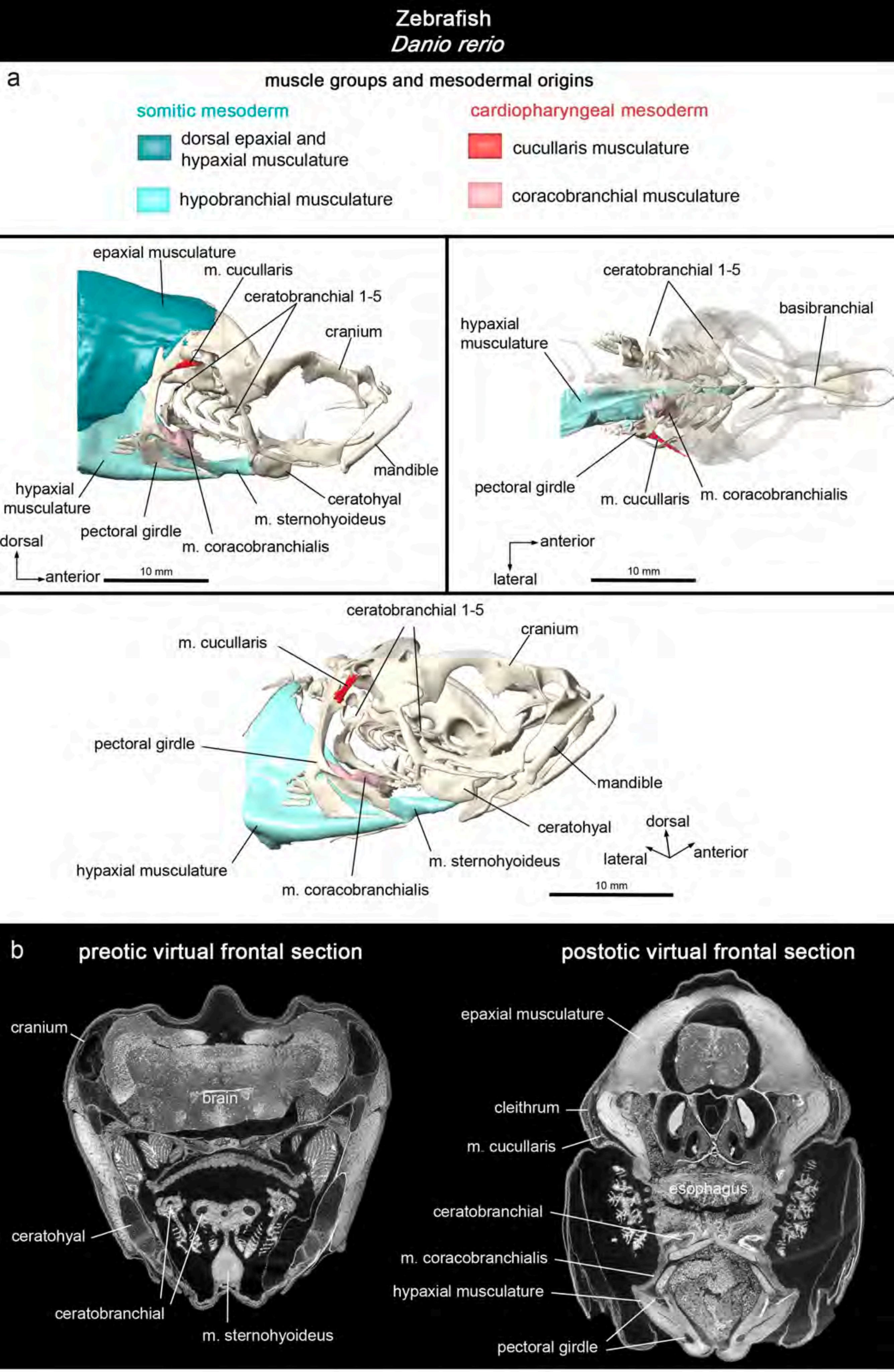


Supplementary Fig. 3

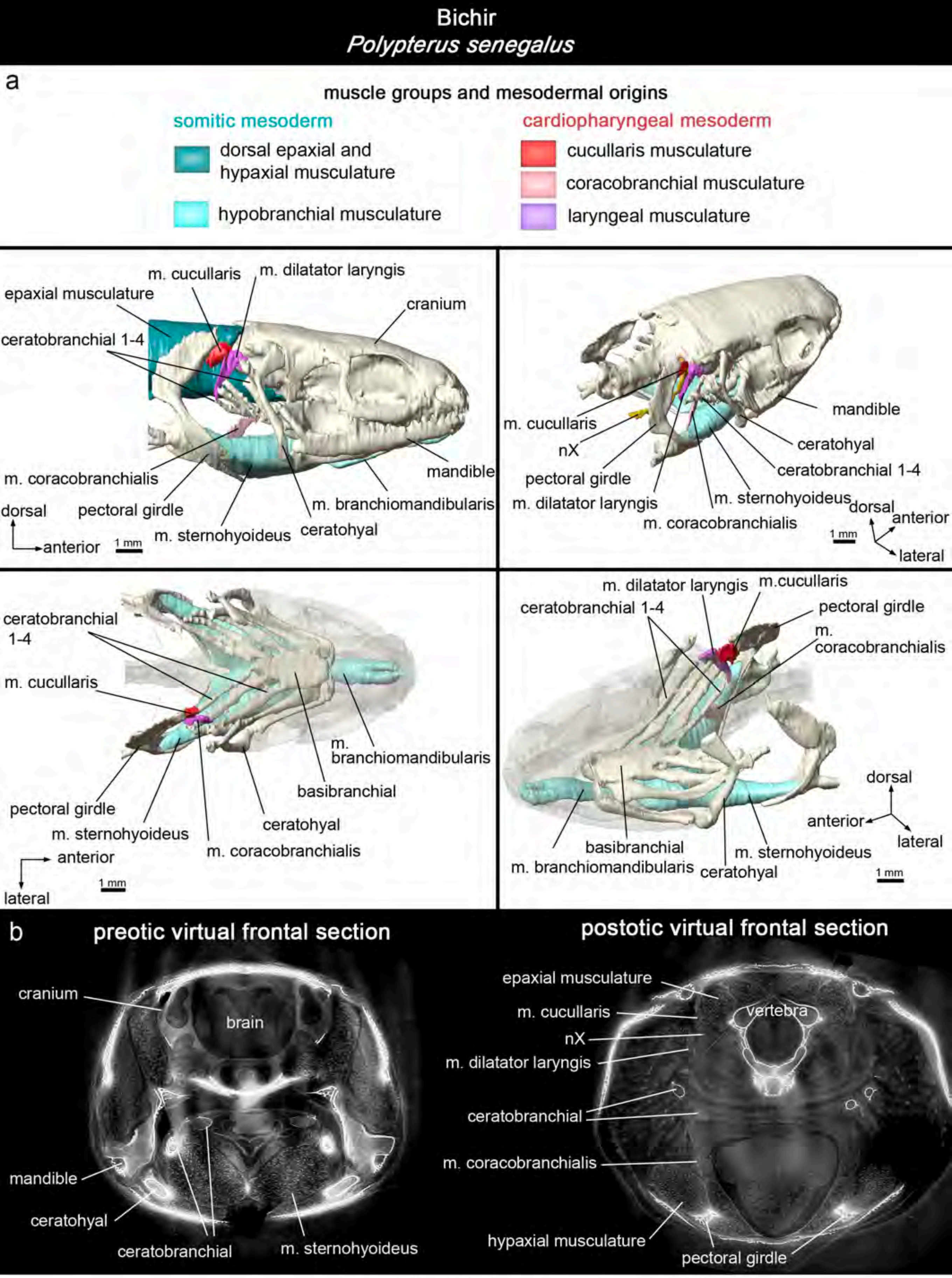


Supplementary Fig. 4

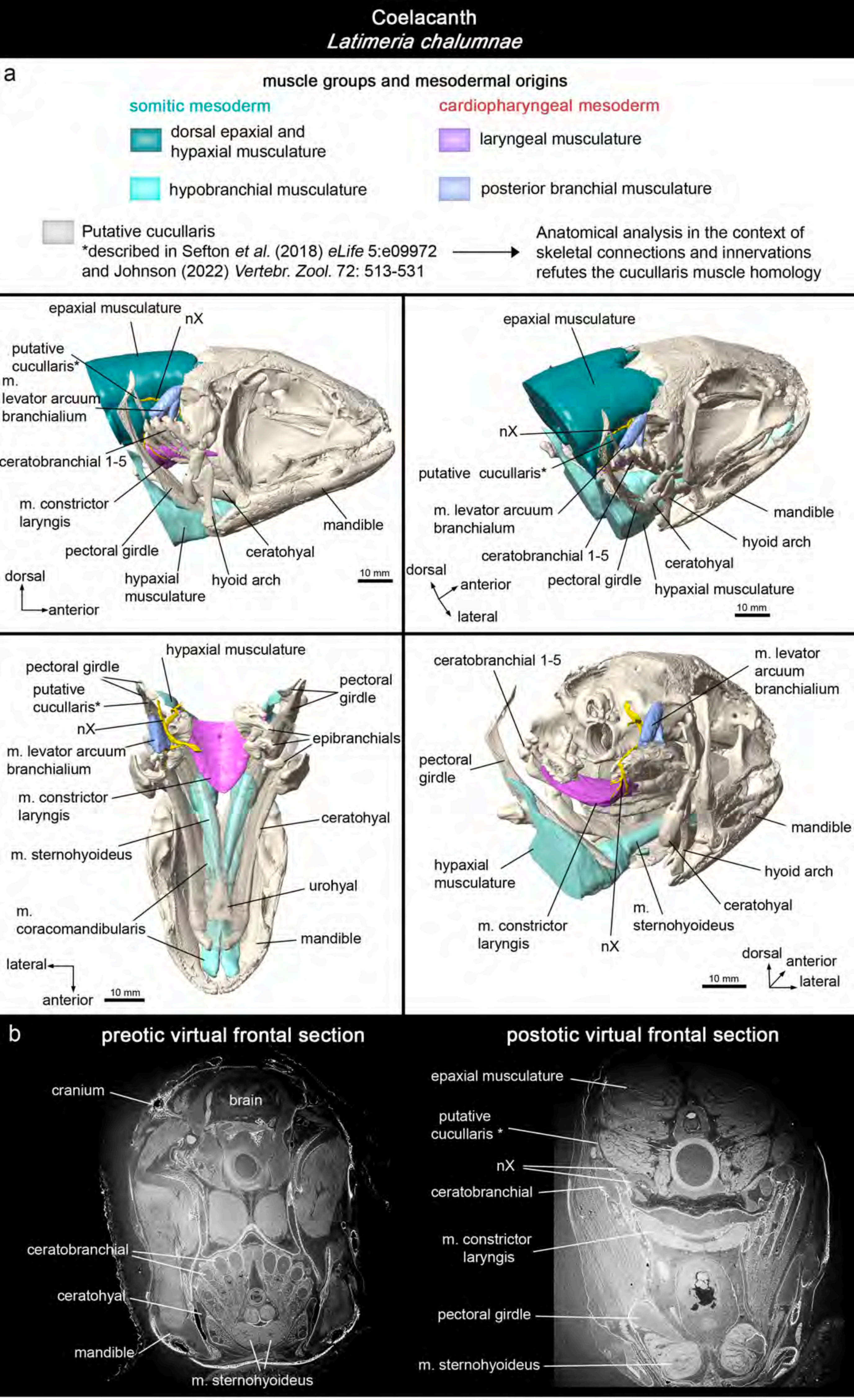


Supplementary Fig. 5

Lungfish
*Protopterus dolloi*

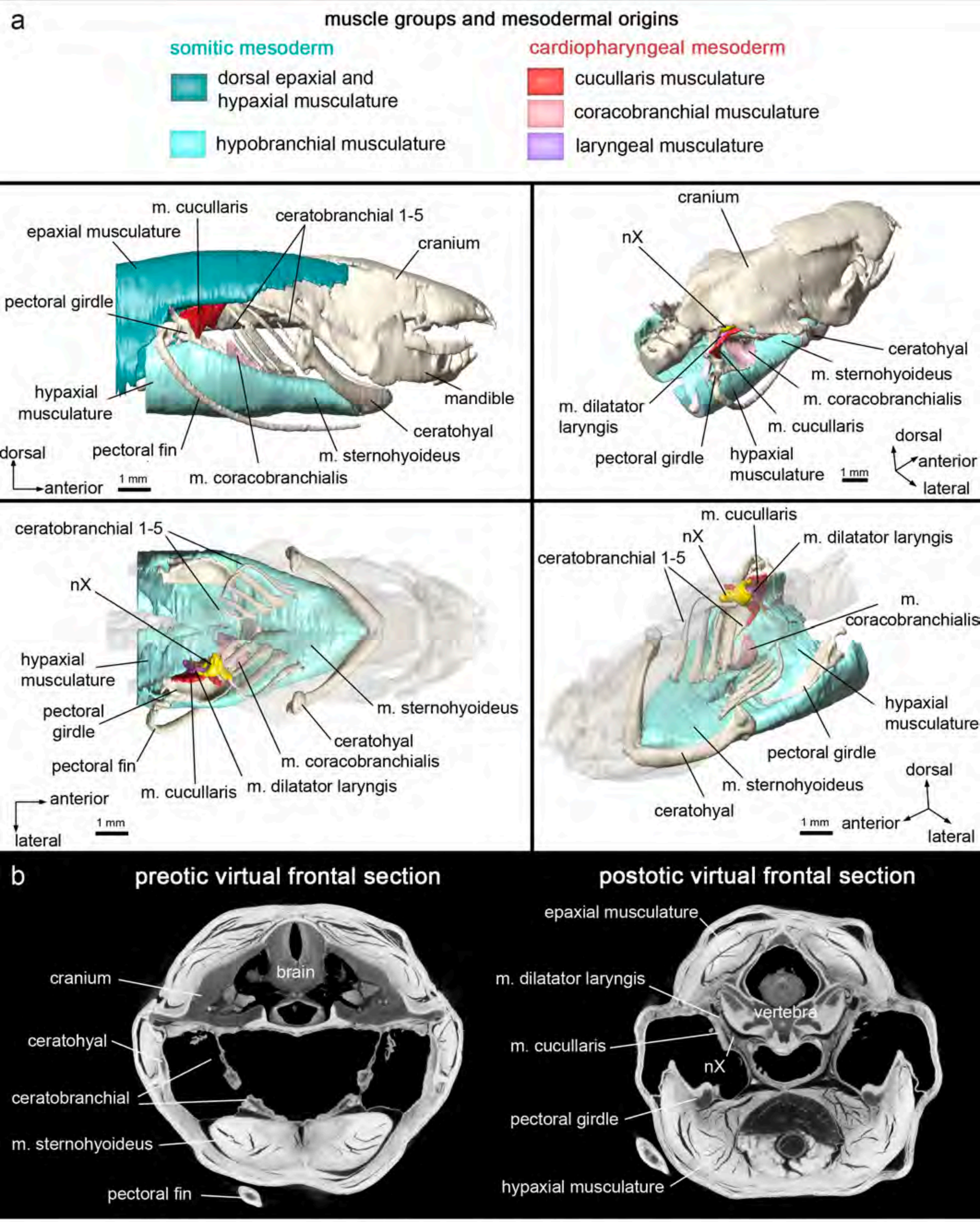


Supplementary Fig. 6

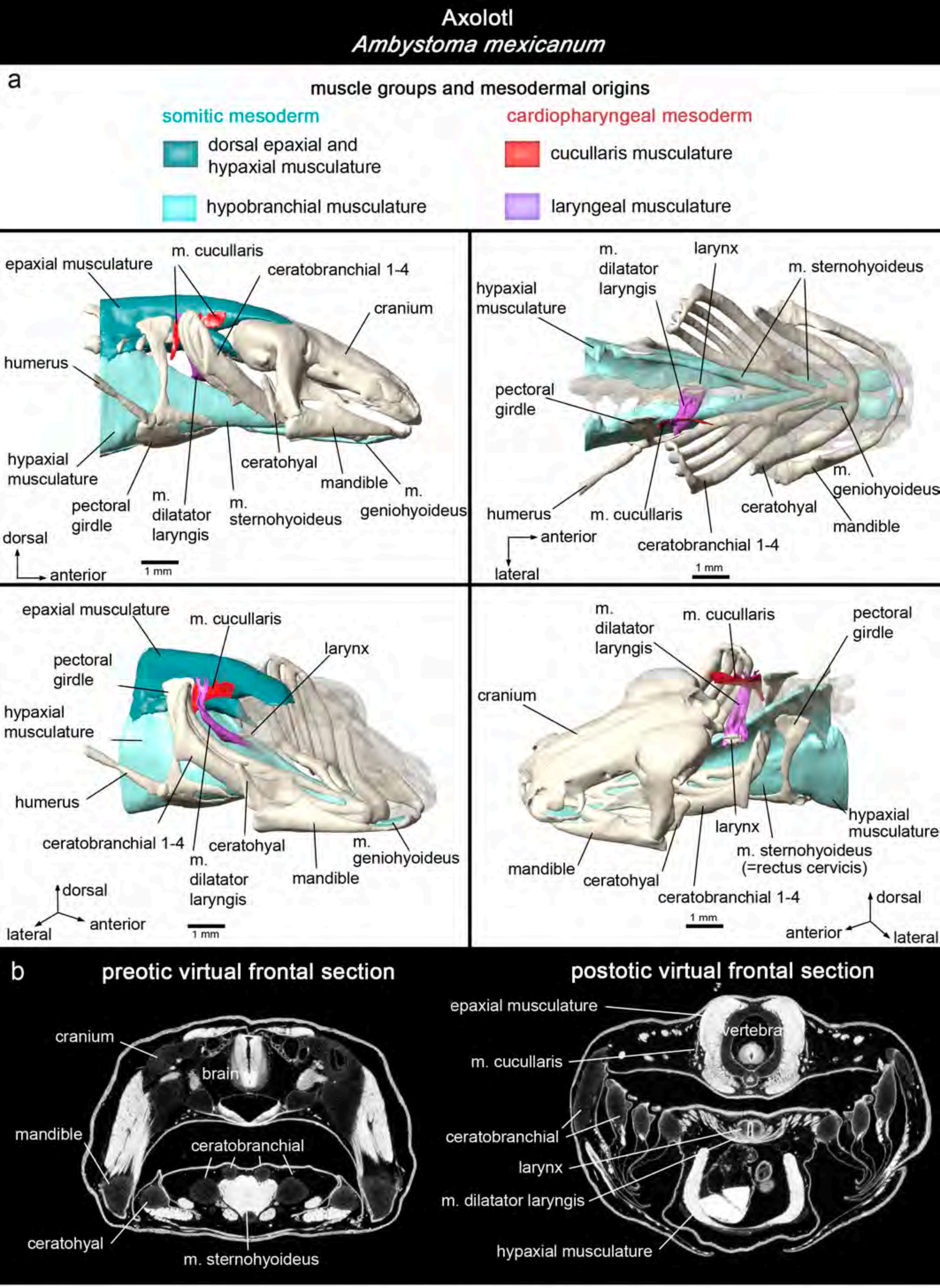


Supplementary Fig. 7

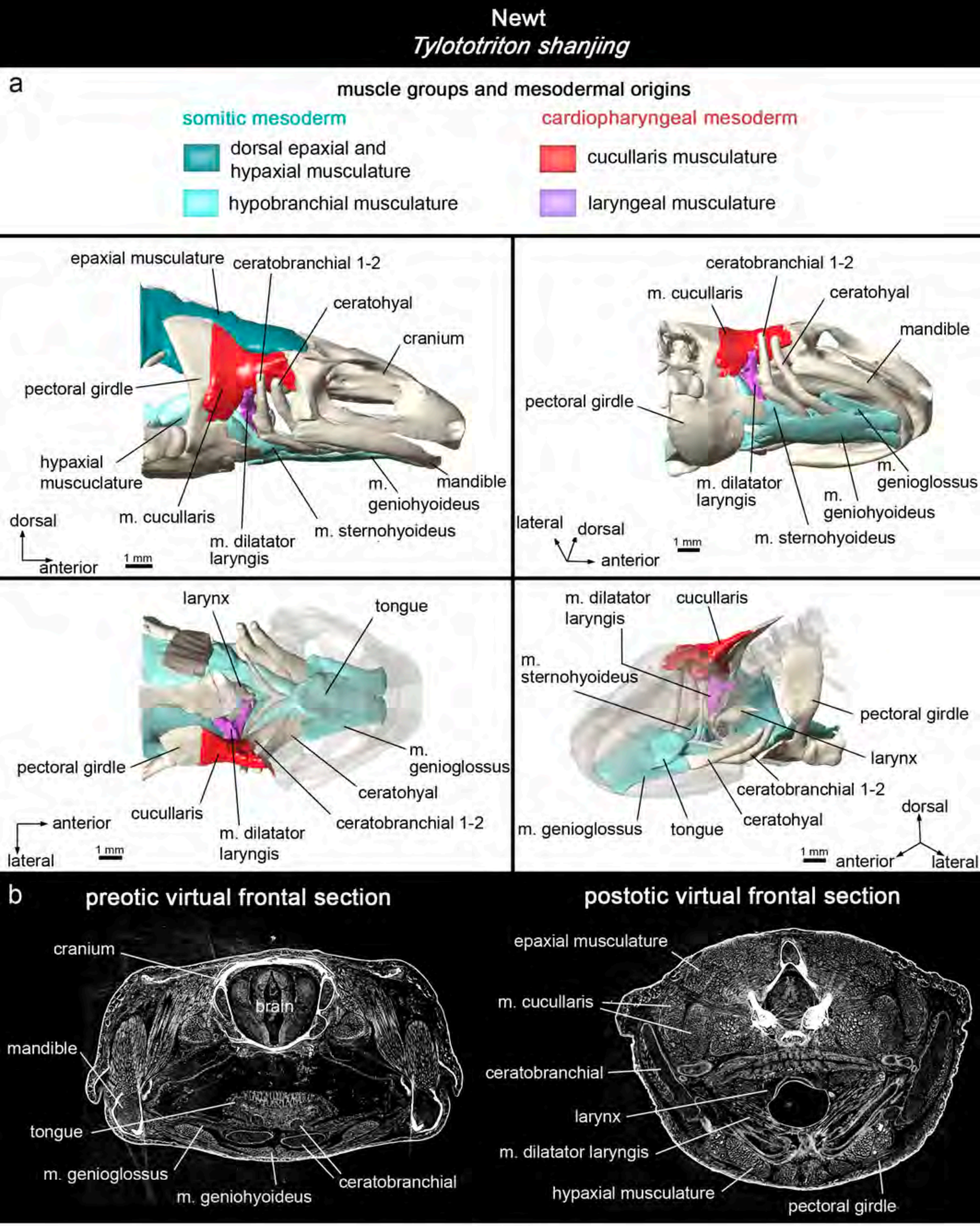


Supplementary Fig. 8

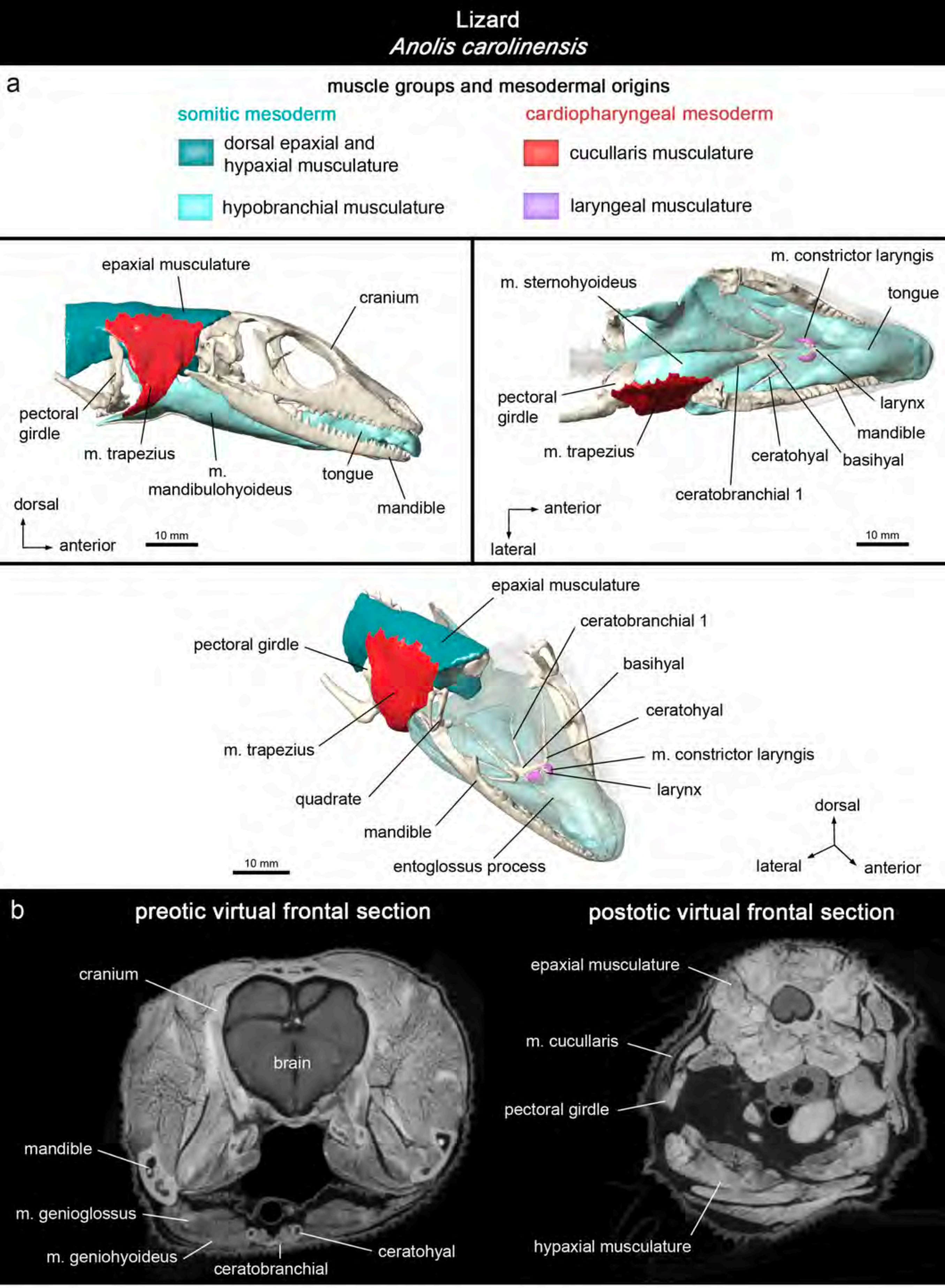


Supplementary Fig. 9
